# Oncogenic *Ppm1d* mutations deregulate the p53 pathway in primary mouse gliomas

**DOI:** 10.1101/2025.11.25.690483

**Authors:** Vennesa Valentine, Abigail J. Groth, Joshua Tolliver, Avneesh Saravanapavan, Spencer M. Maingi, Loren B. Weidenhammer, Nerissa T. Williams, Lixia Luo, Matthew S. Waitkus, Sheeba Jacob, Zachary J. Reitman

## Abstract

**Importance of Study:** Protein phosphatase magnesium-dependent 1D (PPM1D) is frequently mutated in diffuse midline gliomas (DMGs). DMGs are rare pediatric brain tumors with limited treatment options. Due to the cancer’s rapid progression, patients usually survive 12-24 months after diagnosis. This underscores the critical need to better understand the molecular mechanisms driving DMGs. This study describes a novel mouse model that provides a powerful platform to investigate PPM1D-driven tumor biology and offers mechanistic insights into disease development and progression. Furthermore, it serves as a valuable preclinical system for evaluating therapeutic strategies and identifying translational opportunities to target Ppm1d-mutant tumors.

**Background:** Diffuse midline gliomas (DMGs) are incurable brain tumors with limited treatment options. Approximately 20% of DMGs harbor truncating mutations in exon 6 of phosphatase *PPM1D*, which stabilize the protein and deregulate p53 signaling. However, the consequences of these mutations for tumor initiation, progression, and therapy remain unclear.

**Methods:** We developed a conditional *Ppm1d-loxP-exon6-loxP-exon6-E518X-tag* mouse allele (*Ppm1d-flex-6*) that enables lineage-, spatial-, and temporal-specific expression of a DMG-derived truncated Ppm1d protein from its endogenous locus in the presence of Cre-recombinase. Ubiquitous activation of mutant *Ppm1d* was modeled using the Meox2-Cre driver, and primary gliomas were modeled using the RCAS/tv-a system to introduce Cre and PDGFB co-drivers into Nestin-positive neural stem cells. Complementary studies were performed in mouse embryonic fibroblasts (MEFs) expressing truncated Ppm1d following Cre recombination.

**Results:** While Meox2-Cre-driven ubiquitous recombination of *Ppm1d-flex-6* produced muted phenotypes, *Ppm1d-flex-6* recombination in Nestin+ neural stem cells accelerated gliomagenesis. Its oncogenic effect was weaker than complete p53 loss, and it did not accelerate tumorigenesis further in p53-null tumors. Single-cell RNA-sequencing revealed that *Ppm1d-flex-6* gliomas adopt more progenitor-like transcriptional states and upregulate p53- and cell cycle associated pathways. In MEFs, Ppm1d-flex-6 enhanced proliferation and shifted transcriptomic programs toward MAPK and PI3K-Akt signaling, while impairing DNA damage responses, including reduced γ-H2AX induction after irradiation. These defects sensitized cells to radiation and decreased clonogenic survival after ionizing radiation and PARP inhibition.

**Conclusions:** *Ppm1d* mutations confer intermediate suppression of the p53 pathway, consistent with the clinical features of *PPM1D*-mutant DMGs and are associated with radiosensitivity and PARP inhibitor vulnerability.

## Introduction

Diffuse midline gliomas (DMGs) represent an uncommon subgroup of malignant glial tumors that predominantly affect children and young adults. DMGs are characterized by point mutations in histone 3 molecules, such as lysine-27-to-methionine mutations in H3.3 gene *H3F3A*,^1,2^ leading to the term DMG K27-altered in the 2021 World Health Organization classification of central nervous system tumors ^3^. DMGs arise in critical midline structures of the brain that are largely inoperable due to their anatomical locations. Radiation therapy remains the primary modality of initial clinical management, and the imipridone ONC201 was recently FDA-approved for recurrent DMG^4^. However, these treatments are not curative, and DMGs have a median survival of less than two years from diagnosis ^5–7^. Thus, there is an urgent need to understand the molecular mechanisms underlying DMG development and to improve the effectiveness of therapeutic interventions.

DMGs are characterized by frequent somatic mutations affecting cell cycle regulatory machinery, which include mutations in the protein phosphatase Mg^2+^/ Mn^2+^-dependent 1D (PPM1D). *PPM1D*, also known as wild type p53 inducible phosphatase (WIP1), is normally induced by the tumor suppressor and transcription factor p53 in response to genotoxic stress causing cell cycle arrest ^8,9^. PPM1D functions in a negative feedback loop, and dephosphorylates p53 and other cell cycle regulatory proteins, including γH2AX and Chk2, returning cells to homeostasis and allowing cell cycle re-entry after cell cycle arrest triggered by genotoxic stress^9^. Approximately 20% of DMGs exhibit mutations in *PPM1D* ^10^. *PPM1D* mutations in DMG are truncating mutations (frameshift, nonsense) in the final exons of the gene that remove a putative C-terminal degron but keep the phosphatase domain intact, resulting in stability throughout the cell cycle ^10^. *PPM1D* truncating mutations occur in distinct disease states from *PPM1D* amplifications, implying that the truncating mutations function in a manner that is more complex than a simple increase in *PPM1D* gene dosage ^10,11^. Interestingly, *PPM1D* mutations occur in a mutually exclusive manner with loss-of-function *TP53* mutations in DMG ^10^,implying overlapping oncogenic functions. However, several clinical and functional observations suggest distinct roles for *PPM1D* and *TP53* mutations in DMG. For instance, *PPM1D*-mutant DMGs have been associated with relatively longer overall survival ^12^, tumor localization within specific brain anatomic locations (e.g. medulla) ^13^, and with improved responses to radiation therapy ^14^. Additionally, functional analyses suggest that *PPM1D* mutations may confer specific vulnerabilities to agents such as PARP inhibitors ^15^, metabolic agents such as NAMPT inhibitors ^16^, or agents that target cell cycle regulators such as MDM2 inhibitors ^17^. Thus, studying *PPM1D* mutations may offer opportunities to better understand DMG tumorigenesis and to identify actionable DMG vulnerabilities.

Given that PPM1D is tightly regulated in terms of its tissue-specific, and cell-cycle specific expression, tools to finely modulate DMG-associated *PPM1D* mutations are needed. Several prior studies relied on systems that overexpress mutant PPM1D ^17,18^. Other prior studies of DMG-derived *PPM1D* mutations leveraged established human patient-derived DMG lines, which cannot inform on the impact of PPM1D mutations on the process of tumorigenesis ^14,15^. Thus, it remains unclear whether phenotypes associated with *PPM1D* perturbations could be artifacts of these experimental systems or would be likely to faithfully model PPM1D-mutant DMG. To address these gaps, tools to precisely introduce DMG-derived mutations into the endogenous *Ppm1d* genetic locus in model organisms are needed. While several mouse alleles to perturb *Ppm1d* have been generated ^17–21^, there remains a critical need for an allele to conditionally express DMG-derived truncating mutations from the endogenous *Ppm1d* locus in specific tissue lineages in a time-dependent manner.

In this study, we developed a conditional mouse *Ppm1d* allele to study the impact of DMG-derived PPM1D truncations on tumor development, progression and treatment response. Using the Cre-LoxP recombinase system and RCAS/tv-a retroviral gene delivery system we generated mice to express oncogenic, truncated Ppm1d from the endogenous *Ppm1d* locus in the presence of Cre recombinase in neural progenitor stem cells. We validated and characterized the mouse model using a range of molecular and histopathologic experimental approaches. We confirmed the functional relevance of this model through comprehensive studies using mouse embryonic fibroblasts (MEFs), including proliferation and clonogenic assays, analyses of DNA and protein expression, and drug response evaluations. These studies provide insights on the impact of *PPM1D* mutations on DMG phenotypes and therapeutic responses and provide a tool for the DMG and modeling communities.

## Materials and Methods

### DF1 Cell Culture and Retrovirus Production

DF1 chicken fibroblast cells were maintained in Dulbecco’s Modified Eagle Medium (DMEM; ATCC) supplemented with 10% fetal bovine serum (FBS; Thermo Fisher Scientific), 1% L-glutamine, amphotericin B, and penicillin-streptomycin (Gibco). Retroviral vectors were generated using RCAS-PDGFB, RCAS-Cre, and RCAS-luciferase constructs ^22^.

### Mouse Brainstem Injections

Neonatal mice (postnatal day 3-5) were anesthetized through brief hypothermia on ice. Using a Hamilton syringe, 1×10^5^ DF1 cells in 1 µl expressing RCAS-PDGFB, RCAS-Cre, and RCAS-luciferase were injected into the brainstem. For all injections, DF1 cells carrying each RCAS construct were mixed in equal ratios prior to administration.

### Imaging of primary gliomas

Anesthetized mice underwent chemical hair removal prior to imaging. Mice were then injected intraperitoneally with 150 mg/kg D-luciferin (Gold Biotechnology). Following injection, mice were anesthetized using 2-3% isoflurane in oxygen within an induction chamber.

Bioluminescence was measured 15 minutes post-injection using the IVIS Lumina III imaging system (PerkinElmer). Imaging was performed twice weekly, beginning 4.5 weeks after tumor initiation, to monitor tumor progression.

### Immunohistochemistry (IHC)

Mouse brain tumor tissues were fixed in formalin, paraffin-embedded, and sectioned for analysis. Hematoxylin and eosin (H&E) staining was performed using standard protocols by HistoWiz following their respective standard operating procedures. IHC staining for Ki67 (Cat #Bethyl IHC-00375), Olig2 (Cat #ab109186), GFAP (Cat #NB300-141) and MYC (Cat #ab32072) was conducted by HistoWiz. Staining intensity and cell counts for Ki67 and Olig2 were quantified by manual counting of positive cells under 40x magnification.

### Cell Lines, Cell Culture and Irradiation

Mouse Embryonic Fibroblasts (MEFs) were isolated from E12.5-E13.5 embryos obtained from pregnant female mice (Ppm1d^flex-6/wt^) using a standard dissection protocol ^23^. Embryonic heads were used for genotyping, and the remaining embryonic tissue was dissociated to establish *Ppm1d*-control and mutant cell lines. Cells were cultured in Dulbecco’s Modified Eagle Medium (DMEM; ATCC) supplemented with 10% fetal bovine serum (FBS; Thermo Fisher Scientific), 1% L-glutamine, amphotericin B and penicillin-streptomycin (Gibco). Cultures were maintained at 37°C in a humidified atmosphere with 5% CO₂. MEF cell lines were immortalized using Simian Vacuolating Virus 40 (SV40) large T-antigen (GenTarget, LVP016-GP) containing a GFP selection marker. Cre-mediated recombination was induced using an adenovirus expressing Cre recombinase (Adeno-Cre; Vector Biolabs). Polymerase Chain Reaction (PCR) confirmed successful recombination only in *Ppm1d*-mutant cell lines after Cre treatment [flex-6 Forward Primer: GGTTGGGAAATGCCTATGA, flex-6 Reverse Primer: GGCGCGCCAAATCGGATCCACCTA]. For radiation experiments, cells were exposed to varying doses of radiation using an Xstrahl CIX3 irradiator.

### Reagents and Antibodies

Small molecule inhibitors and antibodies used in the study are listed in Supplementary Table 2. All compounds and antibodies were handled according to manufacturers’ recommendations.

### Cell Proliferation Assays

To evaluate cell proliferation, cells were seeded at a density of 500 cells per well in 96-well plates and monitored daily over a 5-day period using the CellTiter-Glo assay, according to the manufacturer’s protocol (Promega, Cat# G7571). Data were normalized to untreated controls and analyzed using the student paired t-tests in GraphPad Prism 10, with significance defined as p<0.05.

### Cell Cycle Analysis

Cells were seeded in parallel to allow synchronized treatment. Confluent plates were counted and resuspended at a concentration of 1×10^6^ cells/mL in a 200 µL volume. Cells were stained with propidium iodide and analyzed by flow cytometry using a BD DIVA Sorter instrument. Data was analyzed using the student paired t-test in GraphPad Prism 10, with significance defined as p<0.05.

### Colony Formation Assays

Cells were plated in triplicate at empirically-determined seeding densities and allowed to adhere for six hours prior to irradiation. Following IR exposure, cells were incubated for 1-2 weeks to allow colony formation, with unirradiated plates as controls. Colonies were fixed and stained with a 6% glutaraldehyde and 0.5% crystal violet solution, rinsed, and counted. Surviving fractions were calculated relative to unirradiated controls. Data was analyzed using the student paired t-test in GraphPad Prism 10, with significance defined as p<0.05.

### Immunoblotting

Cells and tissue-based homogenates were washed with PBS buffer (Thermo Fisher Scientific, Cat #10010023) and then lysed with RIPA buffer (Sigma, Cat #R0278-500ML) supplemented with protease and phosphatase inhibitors (Thermo Scientific, Cat #78442) at 4°C for 45-60 minutes. Tissue lysates were sonicated at 35Amps for 5 mins twice. Protein was measured using the Pierce BCA Protein Assay (Thermo Fisher Scientific, Cat #23225) and equal amounts of protein lysates were then prepared, heated on a 95°C heat block for 5 minutes and subjected to SDS-PAGE (10% Cat #4568035, 12% Cat #4568044, or 4-15% Cat #4561083 gels), followed by transfer to PVDF or nitrocellulose membranes. Membranes were blocked with Intercept PBS blocking buffer or 5% BSA. After blocking, membranes were incubated with specific primary antibodies for P53, Phospho-p53 (Ser15), PPM1D, GAPDH, MYC, CHK1, Phospho-CHK1 (T68), H2AX, Gamma H2AX at 4°C overnight. Membranes were washed with 0.5% Tween 20 in PBS before incubation with Licor goat anti-rabbit or HRP conjugated secondary antibody. Proteins were visualized using infrared fluorophore-labeled and imaged with the Odyssey imaging system or BioRad ChemiDoc Imaging System, respectively.

### RNA Sequencing

Total RNA was extracted from control and mutated-*Ppm1d* cell lines using the Quick-RNA™ MiniPrep Kit (Zymo Research). RNA-seq was carried out at Novogene (Sacramento, CA).

Messenger RNA (mRNA) was purified using poly-T oligo-attached magnetic beads, fragmented and used for first and second-strand cDNA synthesis. DTTP (non-strand-specific) and dUTP (strand-specific) libraries were used during second-strand synthesis. Libraries underwent end repair, A-tailing, adapter ligation, size selection, PCR amplification, and purification. Library concentration was assessed by Qubit fluorometric quantification and real-time PCR, and for fragment size distribution using a Bioanalyzer. Libraries were aligned to the mm39 mouse genome and sequenced on Illumina platforms. FASTQ files were processed to generate normalized expression metrics such as read depth, FPKM, and log2fold changes for downstream analysis.

### Single Cell RNA-sequencing (scRNA-seq)

50 micron scrolls of FFPE sagittal sectioned mouse brain containing tumors were confirmed by H&E staining on adjacent 5-micron sections. The GEM-X Flex assay with 3’ scRNA-seq prep type (10X Genomics) was used to generate cDNA libraries from FFPE tissues. 10,000 cells were targeted per sample. Sequencing of cDNA libraries was carried out using an AVITI sequencer (Element Bio). Clustering, cell type annotation, and data analysis from Ppm1d ^wt/wt^; p53 ^fl/wt^ and Ppm1d ^flex-6/wt^; p53 ^fl/wt^ mice (n = 1) were processed in Seurat ^24^, as described previously ^25^. Tumor cells were scored based on the similarity of their RNA-seq profiles to normal brain cell types from a normal brain atlas ^26^ as described previously ^27^.

## Results

### Generation of a *flex-6* allele for conditional expression of oncogenic, truncated Ppm1d from the endogenous locus

To better understand the role of *PPM1D* mutations in DMG tumorigenesis and progression, we generated a transgenic *Ppm1d-loxP-exon6-wt-loxP-exon6-E518X-Tag* mouse model (here called “*Ppm1d-flex-6*”) in which a truncated form of Ppm1d can be conditionally expressed from the endogenous *Ppm1d* locus upon removal of the loxP-exon6-wt-loxP cassette in the presence of Cre recombinase (Supplementary Figure 1). To enable specific detection of truncated Ppm1d protein, we appended a C-terminal 3xMyc tag. We used CRISPR/Cas9 approaches to target our construct to the *Ppm1d* locus in mouse embryonic stem cells (Supplemental Methods). Resultant chimeric mice were bred, and crosses were carried out to remove a Neomycin selection cassette using FlpO recombinase (Figure 1A). To test if organism-wide *Ppm1d* mutation results in specific phenotypes, we crossed the *Ppm1d-flex-6* mice with a strain in which Cre is expressed ubiquitously early in embryonic development, *Meox2-Cre* (Meox2>flex-6) (Figure 1B). PCR analysis of mouse tail DNA confirmed the excision of the loxP-ex6-loxP cassette in the presence of Cre as predicted, with the expected ratio of mice in the resultant litter (Figure 1C). RT-PCR supported specific expression of mutant *Ppm1d* mRNA (Supplementary Figure 2, Table 3). We then assessed the protein expression of wild-type and truncated Ppm1d, Myc-tagged Ppm1d in specific organs, including the testes and brain, for both Meox2>flex-6 and control mice. As expected, we detected the Myc-tagged Ppm1d only in mice with truncated Ppm1d-E518X, confirming the successful recombination of the *Ppm1d-flex-6* allele in Meox2> flex-6 mice (Figure 1D). Thus, we confirmed that our *Ppm1d-flex-6* allele could provide conditional introduction of an epitope-tagged, truncated *Ppm1d*.

**Figure 1.**
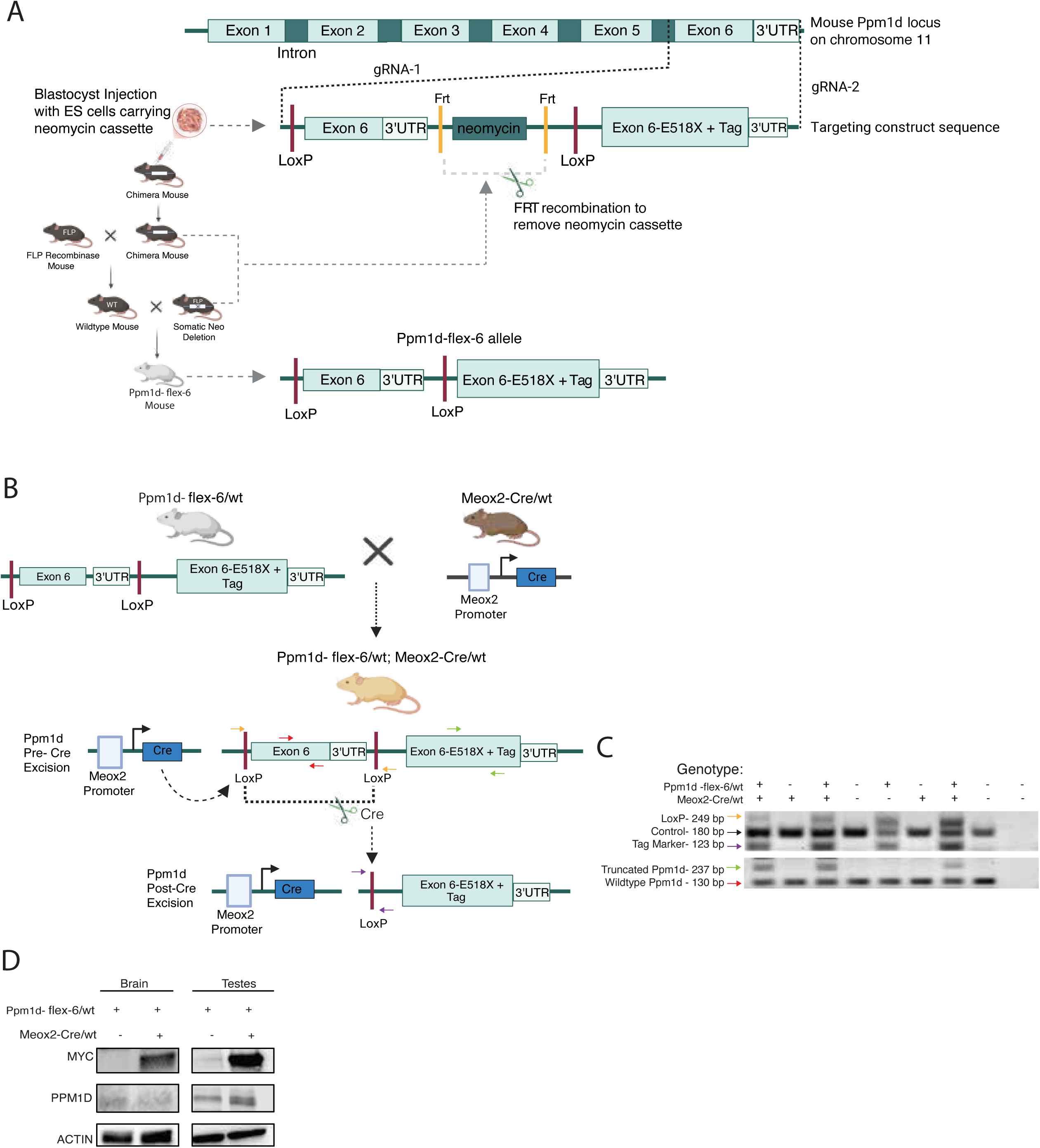
Generation of the *Ppm1d-flex-6* allele and development of the Meox2>flex-6 mouse model. (a) Blastocyst injection with embryonic stem (ES) cells carrying neomycin cassette resulting in chimeric mice. Mice were then crossed with a FLP transgenic mouse to excise the cassette via FLP recombinase, producing a truncated *Ppm1d* allele expressed upon Cre-mediated activation (*Ppm1d-flex-6* mouse model). (b) The *Ppm1d-flex-6* mouse was crossed with a Meox2Cre/wt mouse to generate a model in which the truncated *Ppm1d* allele is activated during early embryonic development through Cre-mediated recombination driven by the *Meox2* promoter. (c) PCR analysis confirms successful recombination at the DNA level. The black arrow represents the actin control. The yellow arrow denotes the unexcised loxP sites in the *Ppm1d* pre-Cre model. The purple arrow indicates the loxP site after Cre-mediated excision confirming recombination and removal of the wild-type exon within the loxP sites. The green arrows denote the truncated *Ppm1d* allele, and the red arrow identifies wild-type exon 6. (d) Homogenates from brain and testes of Meox2Cre>flex-6 mice were analyzed for Myc-tag, Ppm1d and actin by immunoblotting.

### Impact of truncated Ppm1d on mouse development

Truncating *PPM1D* mutations have been associated with cancers including brain tumors and leukemias ^28^, intellectual disability syndromes ^29^, and clonal hematopoiesis ^30,31^. We therefore hypothesized that *Ppm1d* mutation would be associated with specific phenotypes in aged mice. To test this, we observed Meox2>flex-6 mice and compared these to littermate Meox2>*Ppm1d-wildtype* mice over an extended 1-year time period (n=7 mice). None of the mice exhibited gross abnormalities affecting social or developmental behavior. While a trend toward reduced eosinophil levels was observed in mutants compared to controls (p=0.06), no other significant differences were found in overall blood counts or bone marrow profiles (Supplementary Figure 3). One mouse from each group developed lymphoma, but no other tumors were detected. In summary, there was no detectable significant differences between the Meox2>flex-6 and control mice. These findings are consistent with a previous study indicating that mutant *Ppm1d* does not directly affect hematopoiesis in the absence of genotoxic stress ^28^. We conclude that truncated Ppm1d by itself is not a strong driver of tumorigenesis during mammalian development and may require cooperating perturbations to unmask its oncogenic properties.

### Truncated Ppm1d accelerates primary mouse glioma development from Nestin+ neural stem cells

Introducing oncogenic drivers into lineage-restricted stem cells can provide important insights into tumor initiation and development. We hypothesized that truncated Ppm1d expression in neural stem cells, a putative lineage-of-origin for gliomas, in combination with additional oncogenic drivers may accelerate gliomagenesis. To test this, we used the RCAS/tv-a system to introduce Cre and the oncogene platelet derived growth factor beta (PDGFB) into Nestin-positive neural stem cells in mice bearing the *Ppm1d-flex-6* allele and expressing the avian TVA receptor on neural stem cells ^22,32^ (genotype: *Nestin-TVA; Ppm1d-flex-6*) (Figure 2A). We compared littermate *Nestin-TVA* mice with and without the *Ppm1d-flex-6* allele. Truncated *Ppm1d* led to reduced brain tumor-free survival in mice (2-fold increase in tumor-associated mortality at 10 weeks, p<0.05, Figure 2B). The resulting tumors expressed the oligodendrocyte-lineage marker Olig2 and exhibited elevated proliferation, as indicated by the Ki67 marker, while remaining negative for the astrocyte marker GFAP (Figure 2C, 2D). Myc staining was observed in *Ppm1d-flex-6* tumors but not in normal brain tissues or in tumors from littermate controls, confirming tumor-specific expression of truncated *Ppm1d* in the presence of Cre (Figure 2E, 2F). These results show that expression of truncated *Ppm1d* in a lineage and spatially-restricted manner accelerates tumorigenesis in a primary mouse model of DMG.

**Figure 2.**
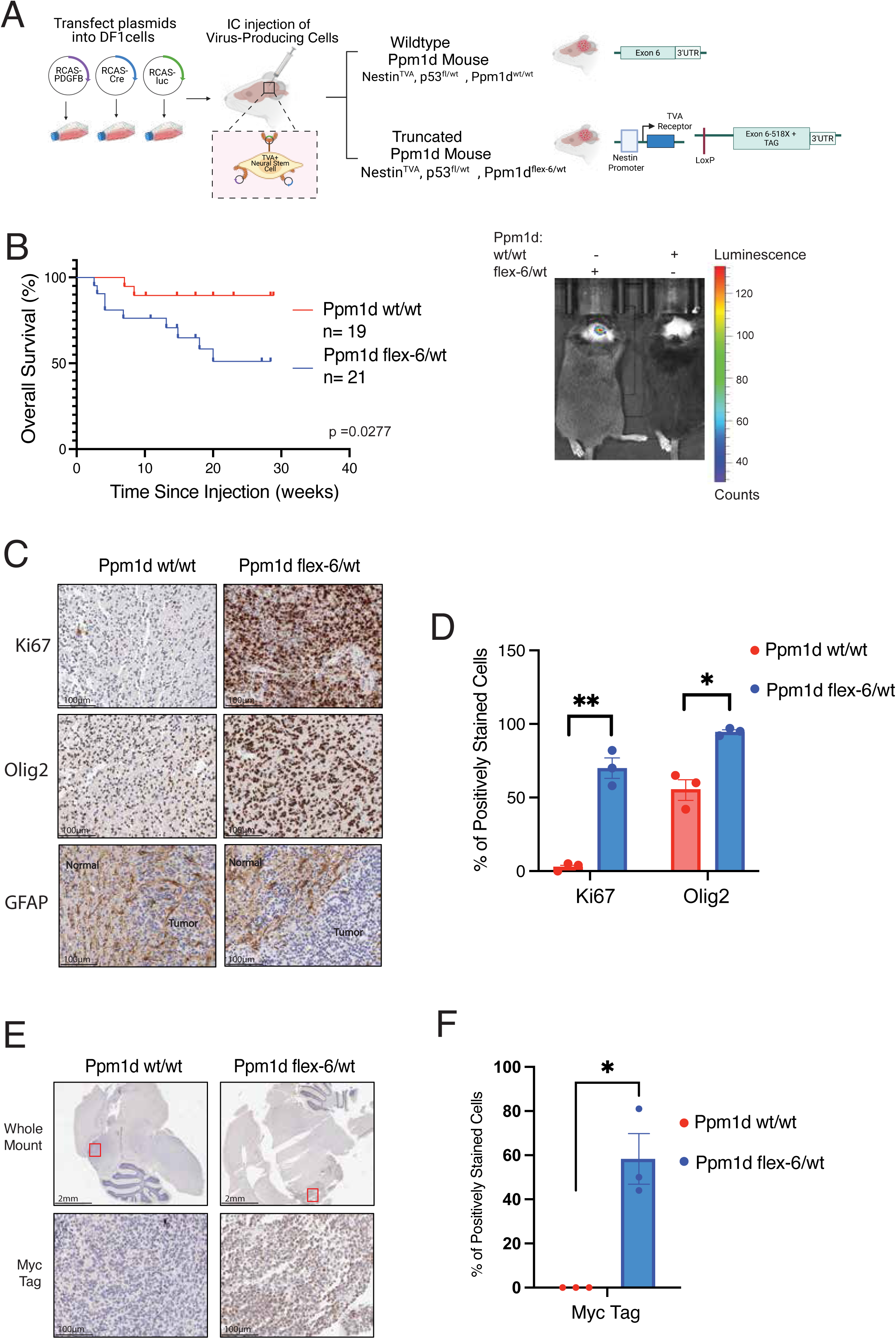
Combining the *Ppm1d-flex-6* allele with the RCAS/tv-a system. a) Intracranial injection of DF1 producing cells carrying RCAS vectors encoding PDGFB, Cre, and luciferase to enable conditional targeting of neural stem cells. b) Mice were injected at postnatal day 3-5 and monitored for tumor development through symptom assessment and bioluminescence imaging starting at week 4 to detect tumor-associated signals. Kaplan-Meier survival estimates and and log-rank (Mantel-Cox) test p= 0.0277, Ppm1d^flex-6/wt^; p53^fl/wt^ n=21, Ppm1d^wt/wt^; p53^fl/wt^ n=19 c) Immunohistochemistry (IHC) with Ki67, Olig2, and GFAP for tumor-bearing brains from wild-type Ppm1d (Ppm1d^wt/wt^; p53^fl/wt^) and truncated Ppm1d (Ppm1d^flex-6/wt^; p53^fl/wt^) mice. Tumor and normal regions were denoted in GFAP staining; GFAP exhibited cytoplasmic localization. Scale bar: 100 μm. d) Quantification of Ki67 and Olig2 staining. GFAP expression was excluded from quantification due to its cytoplasmic distribution. Data are presented as mean ± SEM; n = 3. Student paired t-test, * p< 0.05 and ** p< 0.01 e) IHC staining for Myc-tag was performed on brain sections from wild-type Ppm1d (Ppm1d^wt/wt^; p53^fl/wt^) and truncated Ppm1d (Ppm1d^flex-6/wt^; p53^fl/wt^) mice. Whole-mount sections (scale bar 2 mm) and tumor-region sections (scale bar 100 μm). Red boxes represent tumor regions. f) Quantification of Myc Tag IHC staining. Data are shown as mean ± SEM; n = 3. Student unpaired t-test, ** p< 0.01

### p53 loss drives stronger glioma development than Ppm1d truncation

We next explored effects of mutant *Ppm1d* on the transcriptional cell state of primary mouse gliomas using single cell RNA-seq. After filtering, 10,461 cells were examined from primary glioma-bearing brains derived from Ppm1d-flex-6 or matched littermate Ppm1d-wt mice (Figure 3A). Cells were visualized using UMAP analysis, clustered using k-means clustering, and clusters were annotated based on canonical marker analysis (Figure 3B). Mapping the tumor cells’ transcriptional state to brain developmental hierarchies showed that *Ppm1d*-mutant tumor cells were skewed towards a more progenitor-like phenotype, showing significantly higher mean similarity scores to undifferentiated neurons, but not to dividing neuronal populations, compared to littermate control tumor cells (Figure 3C). Molecular pathway analysis of differentially expressed genes in truncated (*Ppm1d-flex-6/wt)* tumor cells compared to littermate controls (*Ppm1d-wt/wt)* revealed enrichment for TP53-associated transcription of DNA repair genes, chromatin remodeling, and G2/M cell cycle checkpoint pathways (Figure 3D). Together, these findings suggest that the presence of a *Ppm1d* mutation affects the tumor cell state and cell cycle status of tumor cells.

**Figure 3:**
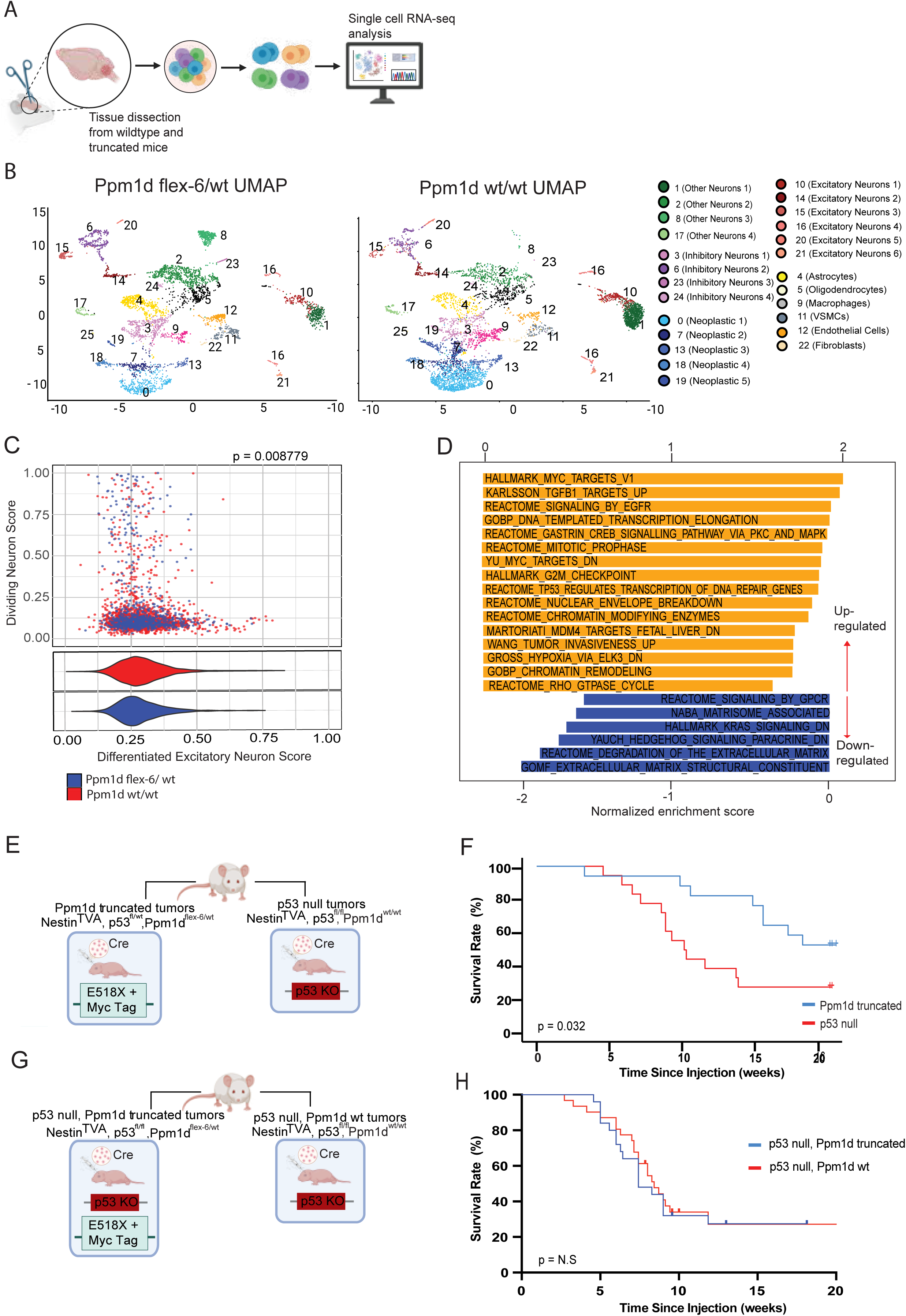
Impact of truncated *Ppm1d* on transcriptomic state and p53 pathway in primary mouse gliomas. (a) Tumor-bearing brains collected from wild-type (Ppm1d ^wt/wt^) and truncated Ppm1d (Ppm1d ^flex-6/wt^) mice were analyzed by single-cell RNA sequencing (scRNA-seq) analysis. (b) scRNA-seq of >10,000 cells from tumor-bearing brains of Ppm1d ^wt/wt^; p53 ^fl/wt^ and Ppm1d ^flex-6/wt^; p53 ^fl/wt^ mice visualized with UMAP plots, with normal and neoplastic clusters annotated. (c) Scatterplots of neoplastic clusters show gene program similarity scores to late excitatory neurons (nEN.late) vs. dividing major ganglionic eminence neurons (MGE.div), colored by condition. Violin plots display nEN.late score distributions stratified by genotype (Ppm1d ^wt/wt^; p53 ^fl/wt^ vs. Ppm1d ^flex-6/wt^; p53 ^fl/wt^). (d) Significant hits from pre-ranked gene set enrichment analysis (GSEA) of neoplastic tumor cells performed using ranked log₂ fold changes between genotypes (Ppm1d ^wt/wt^; p53 ^fl/wt^ vs. Ppm1d ^flex-6/wt^; p53 ^fl/wt^). MSigDB Hallmark, Reactome, and GO collections were tested, retaining pathways at FDR < 0.05. Enrichment results were visualized as bar plots and summarized by normalized enrichment score (NES) direction. (e) Schematic of Ppm1d ^flex-6^; p53 ^fl/wt^ and Ppm1d ^wt/wt^; p53 ^fl/fl^ mice generation. All mice contained Nestin^TVA^. (f) Kaplan-Meier analysis showing tumor-free survival of littermate-controlled mice with either tumoral p53 loss (p53 ^fl/fl^) or tumoral *Ppm1d* truncation (Ppm1d ^flex-6/wt^). Log Rank test P = 0.032, Genotypes: Ppm1d ^flex-6/wt^; p53 ^fl/wt^ n=18, Ppm1d ^wt/wt^; p53 ^fl/fl^ n=22 (g) Schematic of the generated and genotyped mouse models: Ppm1d ^wt/wt^; p53 ^fl/fl^ and Ppm1d ^flex-6/wt^; p53 ^fl/fl^ (h) Kaplan-Meier analysis showing tumor-free survival for littermate mice with tumoral p53 loss and truncated *Ppm1d* (vs. p53 loss alone). Log Rank p= 0.8144, Genotypes: Ppm1d ^flex-6/wt^; p53 ^fl/fl^ n=25, Ppm1d ^wt/wt^; p53 ^fl/fl^ n=11, N.S., not significant.

Given the transcriptomic differences in cell cycle gene program expression, and the links between *Ppm1d* and p53 cell cycle regulatory pathways, we next focused on the impact of *Ppm1d* and p53 mutations on DMG tumorigenesis. Truncating *PPM1D* mutations are mutually exclusive with loss-of-function *TP53* mutations in human DMG, and *TP53* mutations have been associated with shorter overall survival in DMG patients compared to *PPM1D* mutations ^12^.

However, it is unclear whether this clinical association reflects differences in the potency of oncogenic *PPM1D* mutations compared to *TP53* loss, or other clinical factors associated with these tumors. To test whether p53 mutations are a stronger oncogenic driver than *Ppm1d* mutations, we used our *Ppm1d-flex-6* allele to generate littermate mice with primary tumors either expressing truncated *Ppm1d* and an intact p53 allele, or with total loss of p53 (Figure 3E). We found that total p53 loss accelerated tumor formation more potently than truncated *Ppm1d* (4-fold increase in tumor formation at 10 weeks for p53 loss vs. *Ppm1d* truncation, Figure 3F). These data indicate that truncated *Ppm1d* is a less potent driver of DMG than p53 loss.

We next sought to test if truncated *Ppm1d* could promote tumorigenesis independently of p53 loss. To test, this, we examined whether expression from our *Ppm1d-flex-6* allele could decrease time to tumor formation in tumors completely lacking p53. To do this, we generated littermate mice harboring tumors with either truncated or wild-type *Ppm1d* on a p53-deficient background (p53-fl/fl, Figure 3G). Interestingly, survival analysis revealed nearly identical tumor-free survival regardless of *Ppm1d* status in these mice harboring tumors lacking p53 (Figure 3H). These data indicate that *Ppm1d* truncation does not confer additional tumor-promoting phenotypes to gliomas that already lack p53 in mice.

### Truncated *Ppm1d* enhances proliferation in embryonic fibroblasts (MEFs)

Mouse embryonic fibroblasts (MEFs) are useful tools to define the effects of genetic perturbations due to their ease of genetic modification and their physiological relevance ^33^. To explore the oncogenic function of truncated *Ppm1d* in regulating cell proliferation, we derived mouse embryonic fibroblasts from female *Ppm1d-flex-6* mice at 12.5-13.5 days of gestation. MEFs were treated with an adeno-Cre recombinase virus to induce and express mutant *Ppm1d*. PCR analysis confirmed the excision of the loxP-ex6-loxP cassette in Cre-treated cells, with no excision detected in untreated controls (Figure 4A). Immunoblot further validated the presence of the Myc-tagged *Ppm1d* in Cre-treated cells (Figure 4B). We found that *Ppm1d-flex-6* was insufficient to immortalize cells. To facilitate continuous culture of the *Ppm1d*-*flex-6* MEFs, we transduced MEFs with SV40 T-antigen for further experiments (see Materials and Methods).

**Figure 4:**
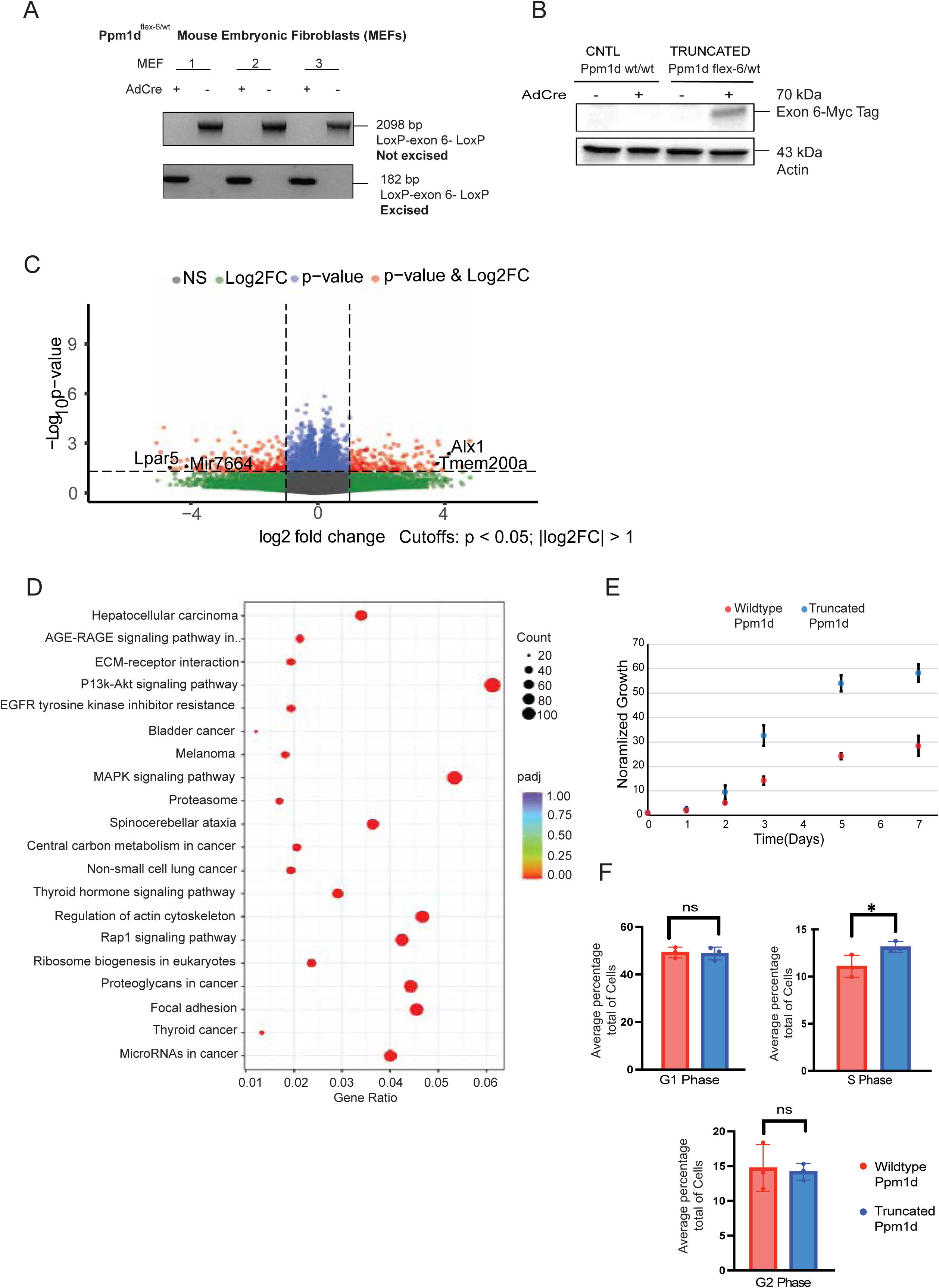
Impact of truncated Ppm1d on the growth of mouse embryonic fibroblasts (MEFs). (a) Three MEFs were isolated from an adult Ppm1d ^flex-6/wt^ mouse at approximately 13 days. PCR analysis confirmed successful removal of the loxP-ex6-loxP fragment in MEFs treated with adeno-Cre (AdCre, 182 bp), whereas non-Cre-treated MEFs retained the full-length loxP-ex6-loxP band (2098 bp). (b) Immunoblot for Myc tag of MEFs with and without AdCre. (c) Volcano plot showing differentially expressed genes by RNA-seq for MEFs with truncated Ppm1d expression (AdCre-treated Ppm1d ^flex-6/wt^) compared to untreated isogenic control MEFs (n=3 biological replicates per group). Cutoffs: p<0.05, log2FC>1 (d) Top significant Gene Ontology (GO) terms from the enrichment analysis. The x-axis denotes the proportion of differentially expressed genes associated with each term, and the y-axis lists the GO terms. Point size represents the number of annotated genes to a specific GO term, and the red-to-purple gradient represents the level of enrichment significance. (e) Proliferation assay in MEF cell lines with Cre-mediated *Ppm1d* truncation (AdCre-treated Ppm1d ^flex-6/wt^) compared to untreated isogenic control MEFs. Relative cell number was measured each day after seeding. Mean from n=4 technical replicates per group. Error bars represent standard deviation. (f) Mean percent of cells in each phase of the cell cycle for MEF cell lines with Cre-mediated *Ppm1d* truncation (AdCre-treated Ppm1d ^flex-6/wt^) compared to untreated isogenic control MEFs. n=3 per group, student paired t-test * p< 0.05. Error bars represents standard deviation.

To characterize the impact of *Ppm1d* truncation on MEFs, we performed RNA-seq on MEF cell lines expressing truncated *Ppm1d* or controls. A total of 1,892 genes were upregulated, and 1,644 genes were downregulated (log2FC≥0, p≤0.05) in cells expressing truncated *Ppm1d* compared to wildtype controls. Top upregulated transcripts included *Tmem200a* and *Alx1*, genes commonly linked to activation of the MAPK and PI3K-Akt oncogenic pathways which are known to be regulated by *PPM1D* ^34,35^. In contrast, *LPAR5* and *miR-766-4* were downregulated, both of which are typically associated with enhanced PI3K pathway activity when expressed at low levels ^36^ (Figure 4C). Pathway analysis confirmed significant upregulation of the MAPK, PI3K-Akt, and Rap1 signaling pathways in the mutant *Ppm1d* group (Figure 4D), which are key regulators of cellular processes such as proliferation, differentiation, and stress response. This enrichment suggests that *Ppm1d* truncation may influence MEF growth by deregulating these signaling cascades. To test effects on cell proliferation rate, we next performed a cell viability assay to assess the impact of *Ppm1d* truncation on cell growth in MEF cells. Truncation of *Ppm1d* conferred a proliferative advantage to the cells over a 7-day period when normalized to wildtype *Ppm1d* MEF growth (2-fold increased growth at day 7, p= 3.6 x 10^-10^, Figure 4E). Flow cytometry was used to examine the basis for this growth advantage and revealed an 19% increase for the proportion of *Ppm1d*-truncated cells in the S phase (p=0.03, Figure 4F). These cells exhibited a numerically shortened G1 phase, indicating accelerated entry into S phase and increased proliferative activity. Moreover, the decreased proportion of cells progressing from S to G2 phase suggests G2/M checkpoint dysregulation. Overall, these data indicate that truncated *Ppm1d* promotes cell growth by stimulating pro-growth pathways and abrogating cell cycle checkpoints.

### Truncated *Ppm1d* increases radiosensitivity and impairs DNA damage responses

Human *PPM1D*-mutant DMGs have been associated with improved responses to radiation therapy compared to *PPM1D*-wildtype DMGs ^12^. We hypothesized that presence of a *Ppm1d* mutation would make cells more vulnerable to ionizing radiation (IR). To test this, we assessed the effect of Ppm1d mutation on clonogenic survival after IR. Cre-treated and untreated MEF cells were monitored for colony formation for 10-14 days following exposure to 0 Gy, 2 Gy, 4 Gy, 6 Gy, and 8 Gy of IR. Notably, Cre-treated cells exhibited reduced clonogenic survival compared to untreated controls at all radiation doses, with a log-fold decrease in colony formation observed at 8 Gy (p=2.5 x 10^-8^, Figure 5A). This finding raises the possibility that the truncation of *Ppm1d* impairs the DNA damage response (DDR), compromising the cells’ ability to recover from genotoxic insults.

**Figure 5:**
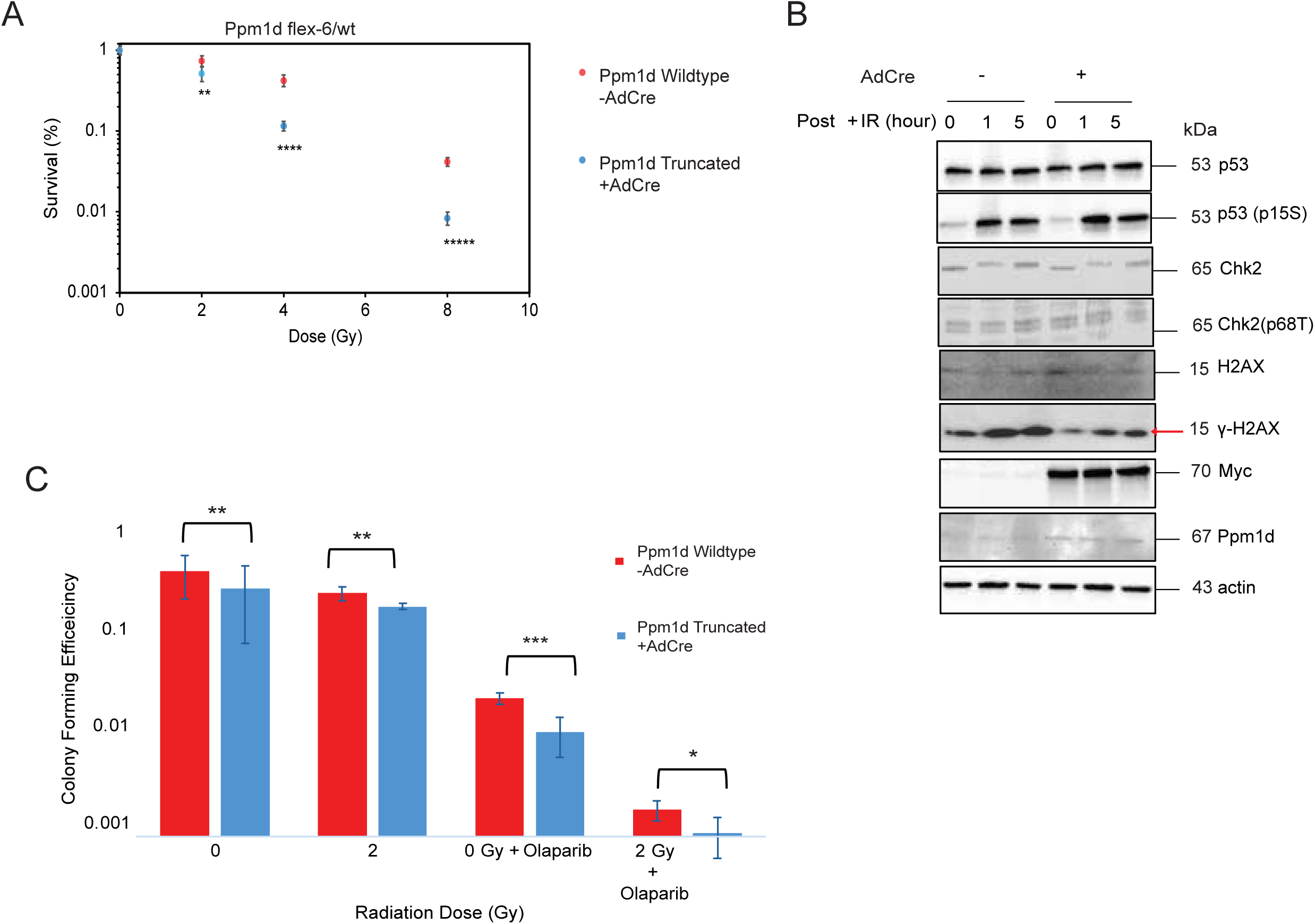
Effect of truncated *Ppm1d* on radiation response and DNA damage signaling. (a) Colony formation in MEF cell lines with Cre-mediated *Ppm1d* truncation (AdCre-treated Ppm1d ^flex-6/wt^) compared to untreated isogenic control MEFs treated with 0 Gy (no radiation) and 2, 4, and 8 Gy ionizing radiation (n=3 per group). Unpaired Student’s t-test; significance denoted as:, p< 0.006 (**), p< 8 x 10^-7^ (****), and p< 2.5 x 10^-8^ (******). Error bars represent standard deviation. (b) Immunoblots from MEF cell lines with Cre-mediated *Ppm1d* truncation (AdCre-treated Ppm1d ^flex-6/wt^) compared to Cre-untreated isogenic control MEFs exposed to 10Gy IR. Cells were collected before IR (0hr) and at 1hr and 5hr post-IR treatment. Whole-cell lysates were analyzed by immunoblot for p53, phospho-p53 (Ser15), Chk2, phospho-Chk2 (Thr68), H2AX, γ-H2AX, Ppm1d, Myc-tag, and Actin. (c) Colony formation in MEF cell lines with and without Cre-mediated *Ppm1d* truncation after 0 Gy (no radiation) and 2Gy ionizing radiation with and without treatment with 1µM olaparib. Error bars represent standard deviation, * p< 0.05, ** p< 0.01, *** p< 0.001

We hypothesized that deregulation of DDR signaling may be associated with the radiosensitivity of *Ppm1d*-mutant MEFs. To investigate the extent to which truncated Ppm1d dephosphorylates key DDR proteins such as p53, H2AX, and Chk2, we evaluated protein phosphorylation status following IR-induced double-stranded DNA breaks. Immunoblots revealed reduced levels of γH2AX in Cre-treated MEF cells compared to controls following 10 Gy IR (Figure 5B). γH2AX reduction was also observed at baseline (Figure 5B). These results show that the truncation of Ppm1d results in decreased phosphorylation of Ppm1d target proteins in the DDR pathway, such as γH2AX, which may underlie increased sensitivity to IR.

### Increased sensitivity to PARP Inhibition after *Ppm1d* truncation

*PPM1D* alterations have been associated with vulnerabilities to several specific agents, including PARP inhibitors ^15^, MDM2 inhibitors ^37^ and inhibitors of PPM1D itself ^14^. To evaluate whether our conditional *Ppm1d*-*flex-6* allele confers drug sensitivities observed in other models, we performed a focused drug mini-screen using MEFs expressing mutant *Ppm1d*. The screen included seven clinically-relevant inhibitors previously associated with sensitivity in *PPM1D*-mutant cells, or proposed for therapeutic repurposing in gliomas ^38^. Although *PPM1D* mutations have been associated with sensitivity to several drugs, our screen did not reveal any inhibitor that selectively reduced the growth of *Ppm1d*-truncated MEFs relative to control cells (Supplementary Figure 4). Notably, both *Ppm1d* mutant and control cells exhibited reduced proliferation after 72 hours of treatment with the PARP inhibitors Talazoparib, Olaparib and Rucaparib (Supplementary Figure 4). The results suggest that *Ppm1d* mutation does not confer vulnerabilities to these seven agents in MEFs, and/or that growth-based screening assays cannot read out *Ppm1d*-mutation-associated vulnerabilities in MEFs.

Since a growth-based readout did not recapitulate drug sensitivities that have been linked to *PPM1D* mutations, we reasoned that a clonogenic readout would be able to recapitulate vulnerabilities to agents that affect replication processes. We hypothesized that PARP inhibition with olaparib could impair the cells’ ability to form colonies when combined with IR. To evaluate this, we performed a clonogenic assay comparing *Ppm1d*-truncated and control MEFs following olaparib exposure at 0 Gy and 2 Gy. *Ppm1d*-truncated cells significantly reduced clonogenic potential relative to controls in the presence of olaparib, IR, or both, with a numerically greater fold-reduction in colony formation observed with the combination (Figure 5C). This observation shows that colony formation assays of *Ppm1d-flex-6* MEFs can be used to read out vulnerabilities that have been associated with PPM1D mutations ^15^.

## Discussion

Here we developed a mouse allele to conditionally express a glioma-derived oncogenic *Ppm1d* mutation from the endogenous mouse *Ppm1d* locus. This represents a critical modeling tool since *Ppm1d* is finely regulated at the transcriptional and protein levels throughout the cell cycle and in specific cell lineages, and since the mutation affects protein stability. Ubiquitous mutant *Ppm1d* expression resulted in muted phenotypes, which may be in line with the fact that humans with germline truncating *PPM1D* mutations develop overall normally without an increased cancer incidence, albeit with an intellectual disability syndrome ^29^. However, mutant *Ppm1d* expression accelerated brainstem gliomagenesis when expressed along with other oncogenic drivers in mouse neural stem cells. *Ppm1d* mutations affected tumor cell proliferation and developmental cell state in these mouse gliomas. Finally, activation of mutant *Ppm1d* using our in MEFs conferred sensitivity to IR and to PARP inhibition based on clonogenic survival assays. These findings suggest that our isogenic MEF model may represent a platform for identifying vulnerabilities associated with *PPM1D* mutations.

Our mouse model provided an opportunity to dissect effects of *Ppm1d* and p53 mutations on primary DMG tumorigenesis ^32^. Interestingly, *Ppm1d* mutations accelerated tumorigenesis to a lesser degree than *p53* loss (Figure 3F), indicating that the *PPM1D* mutation itself may account for the clinical observation of slightly better outcomes for human *PPM1D*-mutant DMGs compared to *TP53*-mutant DMGs ^12^. Interestingly, mutant *Ppm1d* could not accelerate tumorigenesis further when combined with p53 loss, which may explain the rarity of tumors that contain both *PPM1D* mutations and *TP53* mutations, and the observation of several tumors with clonal populations of either one in a mutually exclusive fashion ^39^. Further, we detected increased susceptibility to IR in *Ppm1d*-mutant cells, indicating that *PPM1D* mutations may contribute to the improved responses to radiation therapy seen clinically in DMG ^12^. Thus, our primary *Ppm1d*-mutant DMG mouse model appears to faithfully recapitulate biology that underpins clinical features associated with *PPM1D* mutations in human DMG.

Our study complements a number of mouse modeling tools to perturb *Ppm1d*. These include a *Ppm1d*-deficient model, which revealed postnatal abnormalities following *Ppm1d* knockout ^20^. A Cre-inducible knockout model further linked *Ppm1d* loss in hematopoietic cells to suppressed tumor growth ^19^. In contrast, *Ppm1d* overexpression promoted tumorigenesis, producing cancers similar to those observed in *p53*-deficient mice exposed to low-dose irradiation ^18^, which suggests that *Ppm1d* overexpression may phenocopy partial p53 loss, highlighting a functional interplay between the two pathways in promoting tumorigenesis. More recently, a Cre-inducible leukemia-derived *Ppm1d* allele lacking an epitope tag was used to model hematologic malignancies ^28^. However, none of these systems can express a DMG-derived, truncated *Ppm1d* from the endogenous locus in a conditional manner. Building on these systems, our allele with virally-delivered Cre recombinase allows targeting of the *Ppm1d* mutations to precise developmental timeframes, anatomic brain locations, and cell lineages which may be needed to study brain developmental processes involved in gliomagenesis. The presence of a Myc epitope tag on our construct may allow future advanced proteomic pull-down and imaging studies.

While we focused the current work on DMG, the allele may be useful to study similar truncating *PPM1D* mutations that occur in a variety of human disease states, such as an intellectual disability syndrome ^12^, clonal hematopoiesis ^30,31^, and esophageal dysplasia ^40,41^ which together likely affect several million patients in the US annually.

The current study has several limitations. First, we have so far been unsuccessful in validating parallel mouse strains with additional *Ppm1d* epitope tags such as a fluorophore. Second, we combined *Ppm1d* with the relatively strong oncogene platelet-derived growth factor B (PDGFB) to increase tumor penetrance, but PDGF receptor alpha (PDGFRA) may be a more faithful alteration found in human DMG ^10^. Third, we have not yet combined the *Ppm1d* allele with the H3K27M mutation that frequently co-occurs with PPM1D mutations in DMG, or other co-drivers such as alterations in *ACVR1*, *PIK3CA*, or *ATRX* ^39,42,43^. Future studies will leverage this *Ppm1d-flex-6* allele to explore effects on tumorigenesis, and potential associated therapeutic opportunities, enforced by faithful combinations of mutant *Ppm1d* with other DMG-associated oncogenic drivers.

## Ethics

Animal procedures were approved by the Institutional Animal Care and Use Committee IACUC) at Duke University (Protocol# A114-22-06-26).

## Author contributions

V.V. contributed to experimental design, analyzed data, and prepared the manuscript. V.V. performed molecular and cellular experiments, including flow cytometry, immunoblotting, clonogenic assays, drug treatments, RNA sequencing analysis, and survival studies. A.J.B. performed PCR, RTPCR, clonogenic assays, and survival studies. V.V. and A.J.B. contributed equally to this work and share first authorship. J.T. generated RCAS viruses for in vitro experiments and performed RNA extraction for RNA sequencing. L.B.W., N.T.W., and L.L. assisted with genotyping and mouse breeding. S.J. assisted with immunoblotting experiments and contributed to experimental design. A.S. and S.M.M. analyzed the single-cell RNA sequencing data. M.S.W. provided critical feedback and assisted with experimental design. Z.J.R. designed experiments, supervised the study, and contributed to manuscript preparation.

## Data availability statement

All data generated during this study are included in this published article and its supplementary information files. RNA sequencing data have been deposited in Zenodo with accession doi: 10.5281/zenodo.17429570, and single-cell RNA sequencing data are available on the Broad Institute Single Cell Portal under accession number SCP3337. Schematics were created with BioRender, with Graphical Abstract available at: https://app.biorender.com/illustrations/68e528bd8277fca8c05636c5?slideId=49d41489-61f6-498e-a5e4-778139a4c7d8

## Acknowledgements

We thank David G. Kirsch for critical feedback, and Maria E. Guerra Garcia, Harrison Q. Liu, Gary Kucera, and Cheryl Bosch for assistance with transgenic mouse generation. We also thank Rebecca Bacon at the Duke Pathology Core Facility for performing diagnostic pathological analysis. We thank the Duke Brain Tumor Omics Program and Element Biosciences for assistance with single cell data generation and analysis.

