## Supplementary Figures and Tables for "Oncogenic *Ppm1d* mutations deregulate the p53 pathway in primary mouse gliomas"

Supplementary Appendix:

Supplementary Methods

Supplementary Figures: 4

Supplementary Tables: 3

Supplementary Methods

**Generation of the Ppm1d-flex-6 allele**

All animal studies were performed under protocols approved by the Duke Institutional Animal Care and Use Committee (IACUC). The Ppm1d-loxP-ex6-UTR-loxP-ex6-E518X-3xMyc transgenic mouse allele known as Ppm1d-flex-6 was generated in collaboration with the Duke Transgenic Mouse Facility). Gene targeting of the Ppm1d gene was performed in embryonic stem (ES) cells using CRISPR/Cas9-mediated homology-independent targeted integration as described ^1^. To do this, a pX333 CRISPR/Cas9 activator plasmid containing Cas9 and gRNAs targeting intronic sequences flanking exon 6 of the Ppm1d locus (forward gRNA: 5’- ctcttgtgatgagatgggt-3’ and reverse gRNA: 5’- aagctccggctccattctg-3’) was co-introduced with a mini-circle donor plasmid into mouse ES cells. The donor plasmid contained lox-P sites flanking wild type Ppm1d exon 6 followed by the 3’ untranslated region (UTR). Between the Lox-P sites, a neomycin cassette was inserted between the FLP Recognition Target (FRT) sites. The neomycin cassette served as a positive selection marker. The loxP sites were followed by a second truncated exon 6 with 5’ splice acceptor, with a DMG-derived E518X mutation followed by a 3xMyc epitope tag and 3’ UTR (loxP-ex6-UTR-FRT-Neo-FRT-loxP-ex6-E518X-3xMyc-UTR). After selection and confirmation of cassette integration by PCR, ES cell clones were injected into blastocysts to produce chimeric mice. To remove the neomycin selection cassette, resultant chimeric mice were bred to FLP transgenic mice (Jackson Laboratory #012930) and crosses were carried out to remove the cassette, yielding the Ppm1d-flex-6 mouse allele. To achieve recombination during embryonic development, the flex-6 mice were crossed to Meox2-Cre ubiquitous embryonic Cre driver mice (Jackson Laboratory #026858). Ppm1d-flex-6 mice were also bred to mice carrying the Nestin-TVA allele to achieve lineage specific gene editing ^2^. PCR and immunoblotting confirmed the loxP-ex6-loxP cassette was excised as predicted in the expected mice. Mouse tails were routinely clipped and analyzed by PCR using probes specific to the Ppm1d-flex-6 and Ppm1d-wt alleles at GeneMaster.

**Reverse transcriptase polymerase chain reaction (RT-PCR)**

Total RNA was isolated from Ppm1d control and mutant cell pellets using the Quick-RNA Miniprep Kit (Zymo Research) following the manufacturer’s protocol. Complementary DNA (cDNA) was synthesized from RNA using the iScript cDNA Synthesis Kit (Bio-Rad). PCR amplification was performed using gene-specific primers, and products were resolved on a 3 percent agarose gel containing SYBR Safe DNA Gel Stain (Thermo Fisher Scientific).

**Phenotyping, Necropsy and Histology**

Necropsy, histological evaluation, and complete blood count (CBC) analysis were performed by the Duke Pathology Core Facility. Mice were euthanized, and organs were harvested, weighed and fixed in 10% neutral-buffered formalin, embedded in paraffin, and sectioned for hematoxylin and eosin (H&E) staining at the Duke Biorepository and Precision Pathology Center. Peripheral blood was collected via retro-orbital sinus puncture into EDTA-coated tubes. CBCs were performed to assess hematologic parameters.

**Cell Viability Assays**

Cells were seeded at 500 cells per well in 96-well plates at day 0. After 24 hours, cells were treated with 10 µM of each drug, and viability was assessed 72 hours post-treatment using the CellTiter-Glo Luminescent Cell Assay (Promega), following the manufacturer’s instructions. Data were normalized to untreated controls and analyzed using the student paired t-tests in GraphPad Prism 10, with significance defined as p<0.05.

Supplementary Figure 1. Sequence of loxP-ex6-Frt-Neo-Frt-loxP-

ex6-E518X-Tag-UTR cassette.

A. Schematic representation of the conditional Ppm1d allele construct.
B. 12,127 bp DNA sequence of the Ppm1d allele containing the loxP-ex6-Frt-Neo-Frt-loxP-ex6-E518X-Tag-UTR cassette.

A
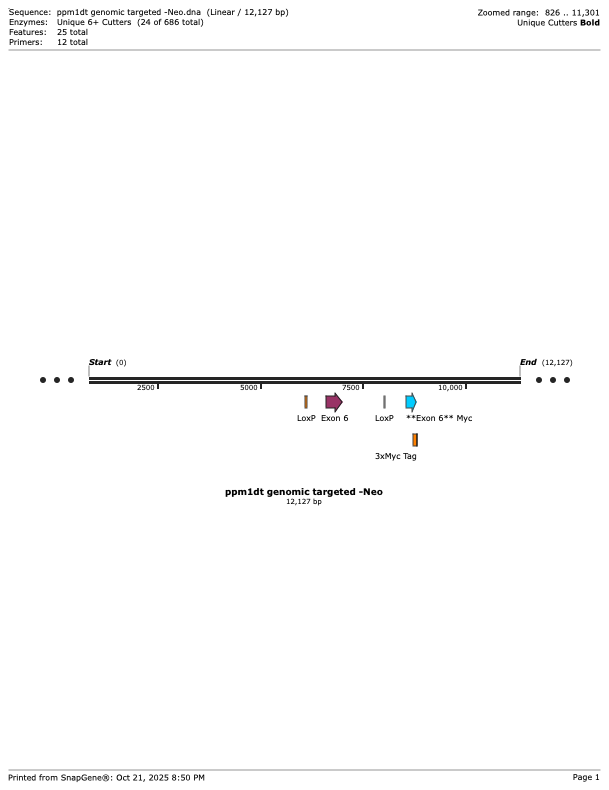

B

gagaatctgaattttattaacttatttggccatactgtagcttttttatttttaaatttttgttttattgtgtttatgcatgcatatggtgtgtatgcgtatgtgttggtcagagggcagtttgaagaggagctggtgactcctatcatgtaagtctagtggattgaactggagaccttgagtcatcaggcttggcagcacccttacctactgagccatctcaccagatctaccaggtctttaatttgaatcatatgcaatggtaagtaattgagagttaatgttcttccttcttgttcttttaaatatagggtgagcaaggacagtcctgtgccaaaatgcttgtgaatcgagcacttggccgctggaggcagcgtatgcttcgggcagataacacaagtgccatcgtaatctgcatctctccagaagtagacaaccaagggaacttcaccaatgaagatgagctctttctgaacctgactgatagccctacttacaacagccaggagacctgtgtgatgacttcttctccaagttctacaccaccaatcaaggtacaaagtactacaggggtaggggtgtgtgtgtgtgtgtgtgtgtgtgtgtgtgtgtgtgtgtgcgtatgtgtgtggtttatttttctattcctgataaagcaccatgaccaagactaacttatacaagaaagtgtttactttgttaatctggcttgtggtttcagagggtcagagtctatgaggacagtgaaagaacagctgagatatttctctctctctctctctctctctctctctctctctctctctctctctaagatttcattttatgtgtatgagtgttttgcttgaatgtaagcctgcttaccatgtgcatgcctggtgctggttgaggttaggagtgaatcccctggagctggaaatatggacggttgtgaactgccatgtggtattgggtcgtgaacctggttcctctgcaagcacaagtactcttaactgctgagccattttgccagccctgagagctcatatcttgatccaaaactgagagagagagaagggggggaggggagagagagcgagcgagtgtgcactgagagaggggggagagagagagagcaagcgagagcgagcgtgcactgggagtctcaagtcttttgaaacctcaaagcctactctcagatacacacctccctcaacaaaggccactcctcctaattctttgccaacagttgtaccagttagggagcaagtgttcagatgtctggggagcagggttgttctcattcatcccagcacatcttgtttgcttttgagatgaggtctcatggtgtggtccatggtgacaaggatgatcttgaattcctgatctttctacttttatctcccacatgttgagattacagtgtttacagtcataattggttaattaagacagaattttggatttgaattaagttcaagaaaatgatggatctttgctctataattcattgctggaaagaatccttttcccctctgcccagataaaccatattctcagttcttcaaagatccatcttcacacgggacctatctgaagtcaggtgtatggatataacttaatagaaaaattattttggccactgaaaactgttaatatgtgaaacttattgatcccttttatgaatctatggacatttgtaaaacatttataagtctgtgaaccacgtgtatacataatagttcatttaaacctataaaacagagctatcctgataggatccatgttatcaataacgacactgaaagattaaatattttgagccaagtcatacagaattaagattgaactggcccttaggatattggaacttagacatgtttttttttttttttttttttttttttaattagcatgggtaagaaacagcattatcttcttatttgacacacgatttcaggcctccaacctgatcctcctatcagagtgccaggattacagaagcatgccaccatgcctgactattcaatttcttagtgtaaatgaaaatggaatgtgtttgagttgtgctagagagagacagctcactggttacaagtacttatctgaaaaacttcagaggacttgagtttggttcccagcatccataccaggcagcacaaaaccatgtgtgactctagcacaagtgggtctgattccctcttctggattcctagggtatccccacacacatagacacacatatatacaaacaaattaaatcttttataaaaaatgtaagctgaggaacacacttaagtgagtacagcatttctgtttcctgatttatttagaactcagttaacctggctttgttaagagtttattttttaggagactggcaattaaagtaggaaataaagcattatgctacatcatcaattaagcaaattttcaaagggccagtgaactgttttgcttttagacaggctgtcatgtagcccaggcaagccttgactcccgggatacaggcctgttctagacaaaggctttggaaggacttttattttggatactgtggttgtgtaaggcaatgactggaatttatcattactatgccatctcttaattcacataaatggcaaaaatagacatgcaaggaagagaagtgtagtctgtgagcagtaattgacaggtttttttttcctttcttcaccccttggtttagtttgcttgtttttgtactacgatagacctcagggtctaatgaatgaagcataaacactctactaagttatatactagttcattacttttggttagccttgaacttaatgtatagcctaggttcacattgactttgtggttctcttgccttagacacctgagtagctatgatcacaggcttgtttcatagtttgagctgattcatagcattaaagttttctgtgttggttatcaaacccagaaagtttggtcaagccacagagttacaccctcttcccctattctaattttattttctttttgtttttttgagacagtgactttctatatatagttctggctgtcctgaaacttacaatgtcgaccaagctggccttgaactcacggagatgtctctgcctctgcctccaaatgttgattagacatgtgtagcaccatgaactcccagctatgtgagtgtgggtgtgcacgccacagcgttcacatggaggtcagaggaggacgttaggctgttttctcctcttgcttttatgtgggtgctggggattgagctccatttgccagactcatgttggaagcatctttatcttacagattgaacgtctcaacagcccacaggacttaaaataaaatttttgtttatttgatgttagtgttttgaaatagggtcttctgtaccacaggctggcttcaaactctttatatatcagaagctggccgtgacctcatgattgttttgccttcatcttccaagtgctgaagttacaggtatgcacttgtacacttggctttcttaaaagaaactttatgttagacgtttcttttcaagccaggcatggttgtgtgtgcctctctgacagctggggtagaaggaccctggattcaaggctaccttgagctaatagcaagactcttcctcaggaagattgccatgagcctggcttaattagagcatgcaaagctaacagattaatatgggtagctacatatgaagttatagctcagaaaaaaaagacttttcactgaaaagtttaaaataatagtacagaatggtacctgttacagtttcatttatttaaggcttcgaagctctgtacctatttgtgactagtgactatgtgaattacttacttgttccctagtgtgcttatggagttgccacttaggcaaatagagaggatctcttagggttagggatccaggcacgaagtgattcctgctgtaatctcaagatctaggaggcttagacaggaggatcactgcaagtcttagagcagcctggactacagagtgagatactgggtaatattatcaagaccctgtcctagaaagagaagggaggggctgcatatatgtctgatgggtaatgggtgatgtgtaagcaggctcactcatctcctaatccccagaatccatgtgtgagccgggcatacatcaggagcatttgtaatcctagcacacctgtgggagatggcatctagagaaaggggaattcatggaagatcatgggcagcttagccggcatacaccacagacaacaaaaacaaagtgaccttgtctcaaacaagactaggactaacaccctaggttatcctctgaccttcatccatgttatggcattctaatacctgctttcacacatatttgttacatacacacatacccactcagaagtgttgtgcacctgtaatcctagtgcttgggaagctgagatagtaggaatctcaagttcaagggcaatatggagagtctgagataattgtaggctacatggtgaaaccctgtctcaaaaaaagaaggggatctattactattcttactgaagtccctcagtagcccgttttcccagttgatgtctggtgtctaatgttcatggggtggggaattacatctgtctttccattatgacataattgctttactgttgtacttcacagttcctagatgtataatagtcacaaattagaatctaggaaagtgacattttttaaaagaacactcttcacttacagttttaattagtttttgtcaagaagccatttataggactggagatctagagttcaggtgatagagagcttgcttgcccagcatgcacgaagccctgggcttgattccctgtctatataagctgagtattggcctcggtctgtaatctagcactataaagtagaggcaggagagtcaaagctgtgatcatccctagttatatagcaagttcaaagccagcctgggttacatgatcaaatgtttttctaaaaaggccattgagaggcatcgagatagtttgcttgctgcctgattcattcctaggatcttcatgatgtgaggagaggtcatatgtaacttgccccctgacctctgcatatgcatattgttgtgtgtgcactctcacatagacgtttttgaaaattgttaattaacaattttgaattgcatttattttgtgtgtctgtatttgttttcccacacacatgccatggcaggacaataatataagcctgtggttcttaatgcattgaagaggaatttactctttttcaccctaattatgtcatcattttgtcacatgtcatatttcagttaaaattgaaaacgagtttatgtaggtaggcattatttgatttaccagaaagttggtatgtcagcttgtttttcatgggcttatgtcaaaaaatagggattttatgtaaaataattagtgctagttcatcagtagtatattattacacataaaattaaaacacagaatttcagctggcagtggttatatataactttaattttagcactggggagacagaagcaggcagatcttagagtttgaaaccagcctggtctacatagccagattcaggacagccagggctatagtgaaaaattctgtctcattaaaaacaaataatgatgggctggtgagatggctcagtgggtaagagcacccaactgctttccgaaggtctggagttcaaatcccagcaaccacatggtggctcacaaccatccgtaatgagatctgactccctcttctggagtgtctgaagacagctacagtgtacttacatatatataaataaatctaaaaaaaaaaccaaaaaagcaaacaatgagtgagtgaaaaaaaacaatgtagagttttgataaagaaattaaaatatcaaaagcagtaattttgccttattctgcatctgggcacagagtgaatgtcttgatagagtgtgtgcttgtggctattttaataggtcttttttgaagccttaaagtattttccttggtcccagctgtggttgggaaatgcctatgagcccatggtcttctccccttgggtgtgtgatgccctctcttgtgatgagatgggttccttctgtgaatcctttgtaccgactgccttctttgcaatcccccaaacataacttcgtataatgtatgctatacgaagttatccaattccgatcatattcaataacccttcccaacttagggtgaggaagtgagttcgtgccatgcttttgccatctgtttgtttgtgtttgagataaggtttttctctggagccctggctgtcctgaacttgctttgtataccaggctggcctcagactcacagagatccacttgcctctgtctcctgagtgctgggactacaggagtgtgccaccactgcccagctggttttactatcttaacaatgtcttaaaaagaacttactgtctgtgtcagtttattgtgcatagaataaatatttatttttccttccctcccactccccagtcaccggaagaagatgcatggccaaggctgagctctaaggaccatatacctgcccttgttcgcagtaatgccttctcagagaagtttttagaggtcccagctgagatagctagagggaatatccagactgtagtgatgacctcaaaagactcagagacacttgaagaaaattgccccaaagccctgactttaaggattcatgattctttgaataatactctgtcagttggcctcattccaaccaattcaacaaatactatcatggaccaaaaaaacttaaagatgtcaactccaggtcaaatgaaagctcaagaagttgaaagaacccctccagccaattttaaaaggacattagaagaatccaactctggcccccttatgaagaagcaccgacgaaatggcttaagtcgaagtagcggggcccaggcttccagtctccctacagcatcccagcgcaggcactctgtcaaactgaccctgaggcgcagactcaggggccagaggaagatgggaaatcctcttctccaccagcaccggaaaacagtgtgtgtgtgctgagatgggcctgggaagtgggggtctccctacctacgactgagggctttttaaacttggtgcgaagttgaacttttttaaggggataaaataaagagaatacagtttgactttttggaatttaacagttttattttggccttgtacttgcctgtattataatgtgaattttgtagatgtagggaataagttgctgtaaaatgtgtgtaaatttgtatcctttacacaagtttagtctcttactctgacacatagtaattgtgacagcagggctaatgttgaagaaaagtcagaagaatctttaagattttaaaaatgtctttaaagtttttaaaatgcttactacatacttatatacaccccttgtgaagaacacatgactttttaaagaaaattactaagcaaactggaaaagtgaagtattttcatagtgatctgtgctccacttaatgtttcccagggaccattagtgtctttttaaaattacattttatttcacatttcataattcagaagtaaacctttcataggaaaaatactgagctgtgctaatgtagctgattttagtctccttgtcccacttacactatgcagtatctcctaacttcagtgcactctgctagaacagtacatttgctgtatttactgaaatctctggcacagaaggaagtgtgtttgcctcacacaccatttgtcccagaccagtggcattaggccatatattctcttctagtgtttgcttaaaatatgtgaagtttttcttgctatttcaataacaaatggtgctgctaacacccaacatttcctaaattattttctatcatacagttttcattggttatatgagtatgtctacccaataaatcactgaatttatgttgctggcgttttgttggtgactcttcatggtaggtgtcgacggtatcgataagcttgatatcgaattccgaagttcctattctctagaaagtataggaacttcatcagtcaggtacataatataacttcgtataatgtatgctatacgaagttattaggtggatccgatttggcgcgccccttctgtgaatcctttgtaccgactgccttctttgcaatcccccaaaccccaacttagggtgaggaagtgagttcgtgccatgcttttgccatctgtttgtttgtgtttgagataaggtttttctctggagccctggctgtcctgaacttgctttgtataccaggctggcctcagactcacagagatccacttgcctctgtctcctgagtgctgggactacaggagtgtgccaccactgcccagctggttttactatcttaacaatgtcttaaaaagaacttactgtctgtgtcagtttattgtgcatagaataaatatttatttttccttccctcccactccccagtcaccggaagaagatgcatggccaaggctgagctctaaggaccatatacctgcccttgttcgcagtaatgccttctcagagaagtttttagaggtcccagctgagatagctagagggaatatccagactgtagtgatgacctcaaaagactcagagacacttgaagaaaattgccccaaagccctgactttaaggattcatgattctttgaataatactctgtcagttggcctcattccaaccaattcaacaaatactatcatggaccaaaaaaacttaaagatgtcaactccaggtcaaatgaaagctcaagagcagaagctgatcagcgaggaggacctggagcagaagctgatcagcgaggaggacctggagcagaagctgatcagcgaggaggacctgtaggaagttgaaagaacccctccagccaattttaaaaggacattagaagaatccaactctggcccccttatgaagaagcaccgacgaaatggcttaagtcgaagtagcggggcccaggcttccagtctccctacagcatcccagcgcaggcactctgtcaaactgaccctgaggcgcagactcaggggccagaggaagatgggaaatcctcttctccaccagcaccggaaaacagtgtgtgtgtgctgagatgggcctgggaagtgggggtctccctacctacgactgagggctttttaaacttggtgcgaagttgaacttttttaaggggataaaataaagagaatacagtttgactttttggaatttaacagttttattttggccttgtacttgcctgtattataatgtgaattttgtagatgtagggaataagttgctgtaaaatgtgtgtaaatttgtatcctttacacaagtttagtctcttactctgacacatagtaattgtgacagcagggctaatgttgaagaaaagtcagaagaatctttaagattttaaaaatgtctttaaagtttttaaaatgcttactacatacttatatacaccccttgtgaagaacacatgactttttaaagaaaattactaagcaaactggaaaagtgaagtattttcatagtgatctgtgctccacttaatgtttcccagggaccattagtgtctttttaaaattacattttatttcacatttcataattcagaagtaaacctttcataggaaaaatactgagctgtgctaatgtagctgattttagtctccttgtcccacttacactatgcagtatctcctaacttcagtgcactctgctagaacagtacatttgctgtatttactgaaatctctggcacagaaggaagtgtgtttgcctcacacaccatttgtcccagaccagtggcattaggccatatattctcttctagtgtttgcttaaaatatgtgaagtttttcttgctatttcaataacaaatggtgctgctaacacccaacatttcctaaattattttctatcatacagttttcattggttatatgagtatgtctacccaataaatcactgaatttatgttgctggcgttttgttggtgactcttcatggtaggtaccatcagttagatatgtaatgtaactgtgctttcttgacttgaaatacatacagtgctcattctttattttaatacagaacacttagaaacaaactaaatacctaacaacacgattcaggtacagcttctgagtgttcagtaaagctccggctccattctgacgggaaatactgaagcgccgagattctataaatgtgctatgtgtgcagttgtgctcttgatttggaacaagcagcctccttgagggctttgggtgtgcctgagttggaagagtgcgtgccttacacacaggaagccctaggcttgatcccagcattgcataaaccagaggcaggaggatcagggatccaaagtcatccccagttacaaagttttaggtcaacccgagaatcatgagacagggcctcaaaacagtagtggataaaacaaaagcagagagagaaacgaggctcaggagaactaagagaaggactagtgggagggtttgctcaacaggtgcaaggccctgtatttaaaaatatagttaggggttggggcgttagcttgttggtaaagtgcttaaggccctgggtttggtcctcagctctggtgaagcagggtgagggaagagaaattcattagtattttatgtgtatgatcattttgcttgcatctaagtacatgtactacgtatatacttagtgagcttagaggcctaagagggtgtcagatctcctggaactggatttaaaggcaaggttttttttttgttgttgttttgttttttgtttttttggttttttgttttgttttgtgttttttttttttttttttttttttttttgatttttcgaaacagggtttctctgtatagccctggctgtcctggatctcactttgtagaccaaactcgaactcagaaatctgcttgcctctgcctcccaagtactgggattaaaggcatgtgccaccacaccaggcttttttttttttaaagtaatttttttacatgtatttatttattatttttcatggtttattttatatgagtacattgtcgctgtcttcagacattccagaagagggcatcagaacccattacagatggttgtgagccaccatgtggttgctggaaattgaactcaggatctctggtagagcagtcagtgctcttaaccactgagtcatctctccagcctataaaggcaggttgtaaaccagtaagtgggtactggaaatcaaaaagagaccttcaggaagaccagctagtgttcttaacttctgagccatctctctggccataatgcctggtatttgatctgtaaccctagcaaaataaaggctgatttggatatgctccgtggcctactatatttcggtactgactcctgggatggcaccgttgaaatactgaaacctttaggaagtggacccactaggatgactatgggaatgcacttaaagggagttataggacccttctgataattattaaatatgtaaattgtataaattggtttcattgacatgtctcatgaggctaccctgcaactttaggtctatctttcaactgtctttccttctcatatctgcctctgcccagtcctgaatagttcctgagttaaattcttttggttgcctgactttggatctgtaggtttgtattcccccccccccccccagacagggtttctctgtgtagcccctggctgtcctggaactcactttgtagaccaggctggcctcaaactcagaaatccgcctgcctctgcctcccgagtgctgaggccggatttctgggcttaggtttgtattctttaccaaatttaggaatcttttagctattatcaactttttctaacactatcctttttattgattctttccagttgcacaccatgcaccccagtctggctcatctcccaggcccctcatgttcgttcttcgaacttgcaactccccccacagataaaaaacaaacaaaaccccaaaagcatagaaaccatctcaccgtggaagctgtagtatgtcacagcgtgtcccacaggataccactctgtccacatcttcactcgaaagtgttcaatgaatcactggcctggttcgagatctctggcttctgtgacaccatcgatattggattctcattggaaaaaaaaaaaaaaaaa

Supplementary Figure 2. RT-PCR to confirm recombination and expression at RNA level for Ppm1d-flex-6 model.

A. Primers were designed to bind within Ppm1d exons and span the intron 6 junction. While primer sets 1 and 2 bind to the wildtype Ppm1d, primer sets 3 and 4 are specific to the flex-6 allele.
B. The table shows the expected band size for all allele combinations used for each primer set.
C. Gel results electrophoresis results matched the expected band length for each primer pair. Primer set 1 confirmed that splicing for Ppm1d wt/wt is occurring properly, as indicated by the 695 bp band. Similarly, primer set 2 also confirms Ppm1d wt/wt expression at the mRNA level with the 368 bp band. Primer set 3 confirms that splicing for the Ppm1d flex-6/wt is occurring properly, as indicated by the 486 bp band. The faint band in the sample containing a flex-6 allele without Cre expression suggests possible leaky recombination. Primer set 4 confirmed that exon 6 at the mRNA level contains the expected 3x MYC epitope tag.

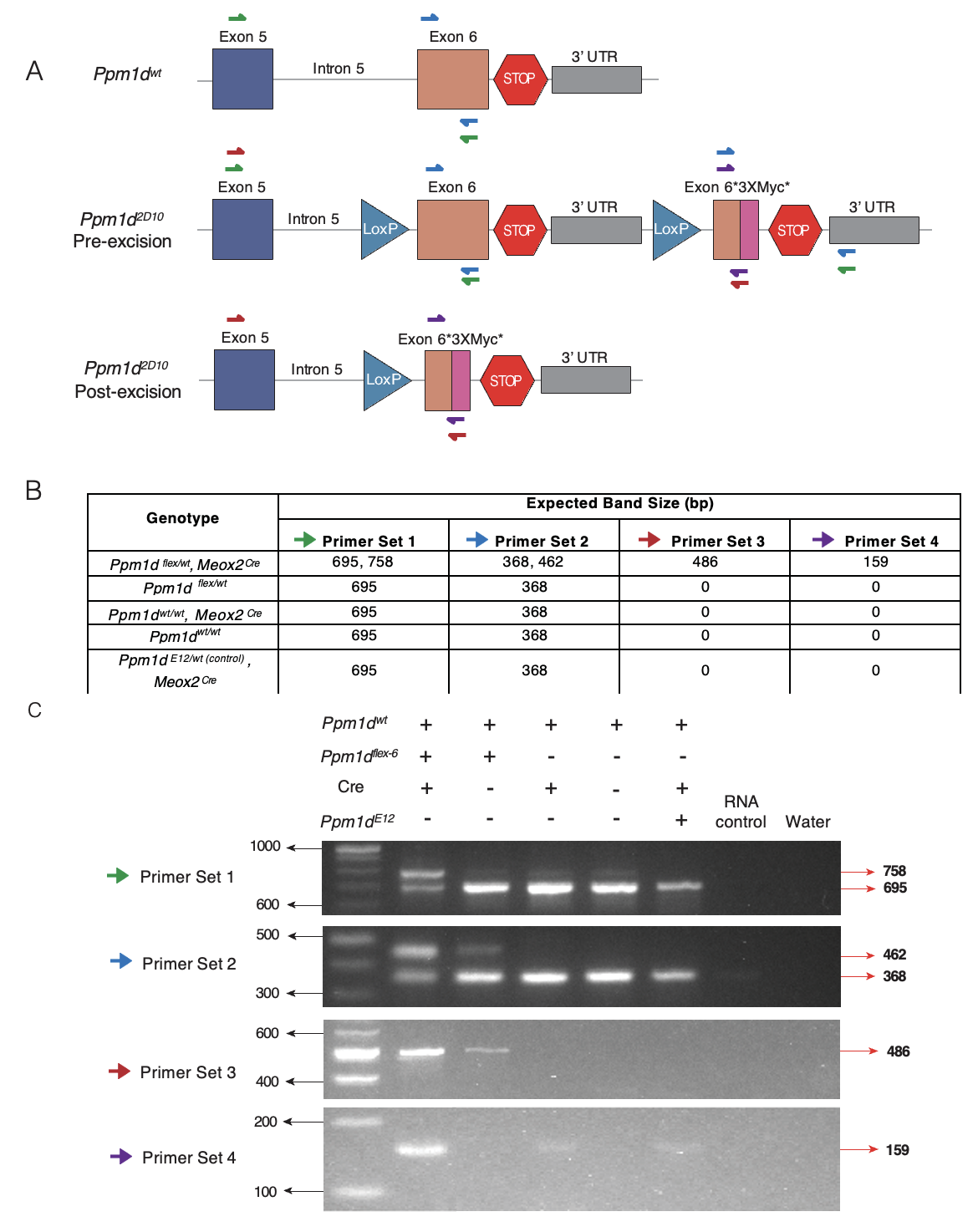

Supplementary Figure 3. Blood counts from Meox2-Cre; flex-6 mice (vs. Meox2-Cre mice)

Peripheral blood was collected from both Ppm1d wildtype and truncated Meox2Cre mice at 1 year for hematological analysis (Meox2Cre; ﬂex-6: n = 4; Meox2-Cre: n = 3). WBC, white blood cells; MONO, mononuclear cells; NEU, neutrophils; EOS, eosinophils; LYM, lymphocytes; BAS, basophils; PLT, platelets; RBC, red blood cells. Error bars represent ± SEM.

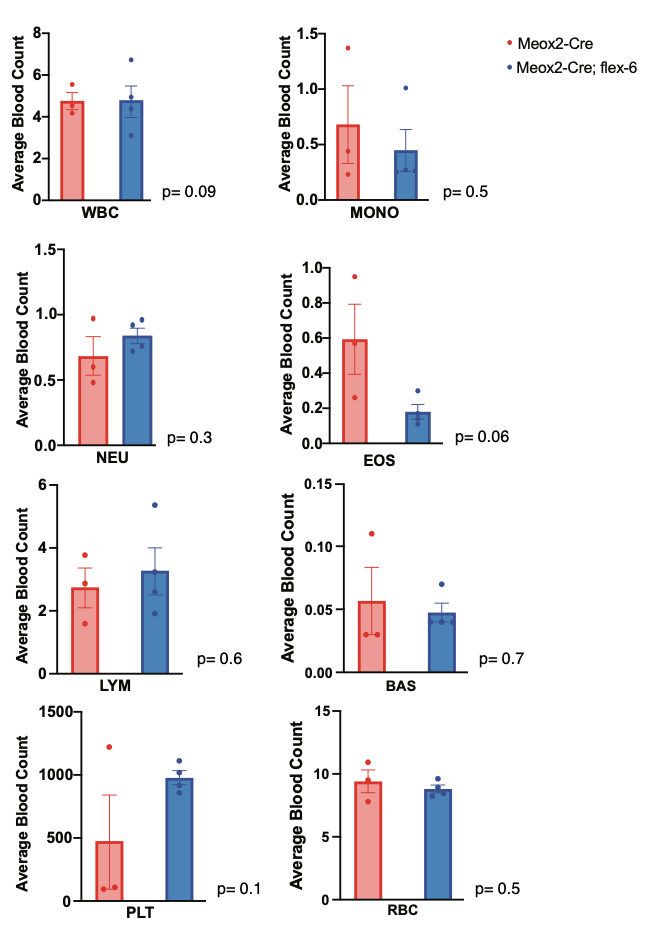

Supplementary Figure 4. Treatment of Ppm1d-flex-6 mouse embryonic fibroblasts (MEFs) with candidate drugs.

MEF cell lines with Cre-mediated Ppm1d truncation and wild-type controls were seeded at 500 cells per well at day 0. After 24 hours, cells were treated with 10 µM of each drug, and viability was assessed 72 h post-treatment. Error bars represent standard deviation.

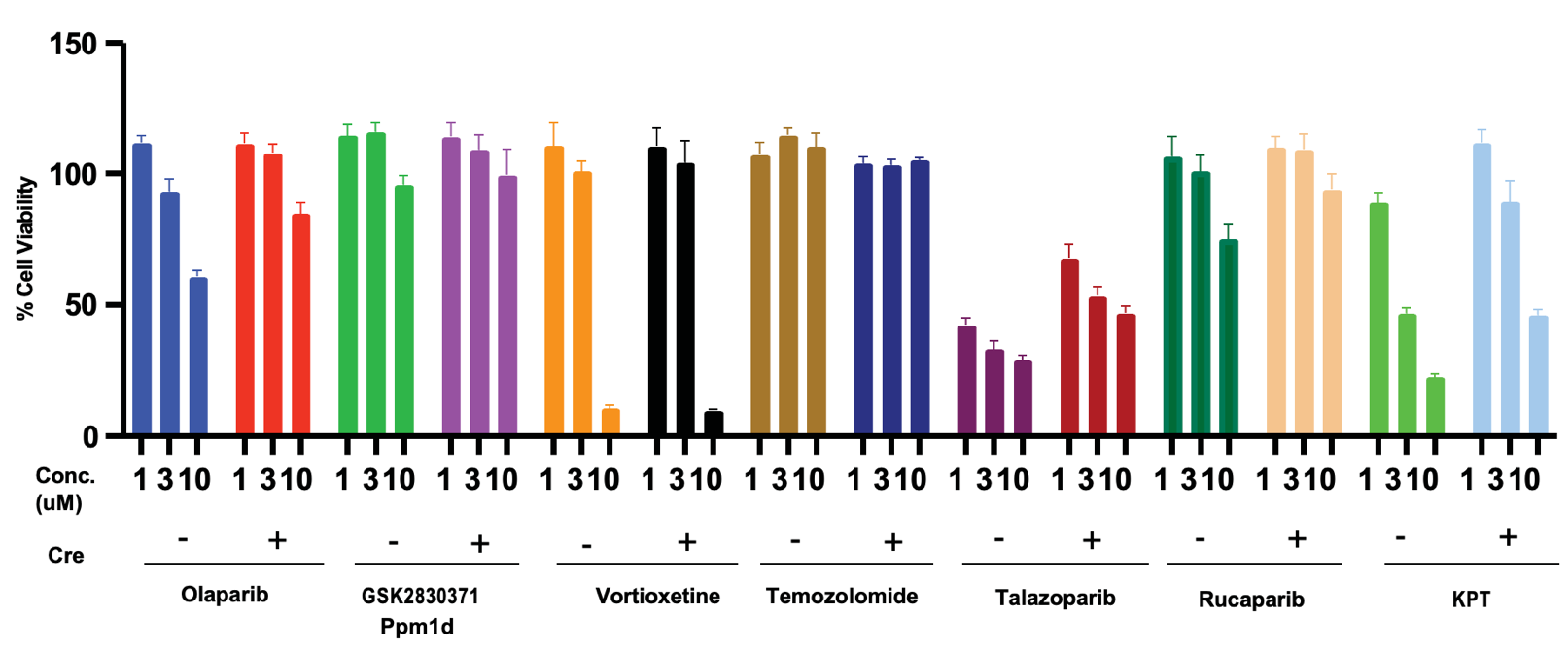

Supplementary Table 1. Probes used for genotyping mouse tails specific to the Ppm1d^flex-6/wt^ allele.

| Primer | Sequence (5' → 3') |
| --- | --- |
| Left | TCTTGTGATGAGATGGGTTCCT |
| Right | GTTGGGGTTTGGGGGATT |

**Supplementary Table 2.** List of small molecule compounds and antibodies used throughout the study. All reagents were handled, stored, and used according to manufacturers’ instructions.

| **Category** | **Reagent / Antibody** | **Supplier** | **Catalog #** |
| --- | --- | --- | --- |
| **Small Molecule Compounds** | Vortioxetine | RayBiotech | 331-10308-2 |
|  | Temozolomide | TargetMOI | T1178 |
|  | Olaparib (10 mM stock) | MedChemExpress | HY-10162 |
|  | Rucaparib Phosphate | MedChemExpress | HY-10617 |
|  | GSK2830371 (WIP1 inhibitor) | Focus Biomolecules | 10-4180 |
|  | KPT-9274 | Fisher Scientific | 50-194-8232 |
|  | Talazoparib | MedChemExpress | HY-16106 |
| **Primary Antibodies** | p53 (D2H9O) rabbit mAb | Cell Signaling Technology | 32532 |
|  | Phospho-p53 (Ser15) (D4S1H) rabbit mAb | Cell Signaling Technology | 12571 |
|  | Histone H2AX (D17A3) rabbit mAb | Cell Signaling Technology | 7631 |
|  | Phospho-Chk2 (T68) rabbit mAb | Abcam |  |
|  | Myc tag rabbit antibody | Abcam | ab9106 |
|  | WIP1 (D4F7) rabbit antibody | Cell Signaling Technology |  |
|  | Chk2 rabbit antibody | Cell Signaling Technology | 2662 |
|  | β-Actin (13E5) rabbit antibody | Cell Signaling Technology |  |
|  | γH2AX (phospho-S139) antibody | Abcam |  |
| **Secondary Antibodies** | IRDye Goat anti-Rabbit | LI-COR | Lot #D20333-15 |
|  | Anti-Rabbit IgG, HRP-linked | Cell Signaling Technology | 7074P2 |

Supplementary Table 3. RT-PCR Primers were designed using the NIH primer design tool and cross-referenced with the mouse genome in the UCSC Genome Browser.

| 1 | Exon 5 F | TGCTTCGGGCAGATAACACA |
| --- | --- | --- |
|  | Exon 6 R | GTGCTGGTGGAGAAGAGGAT |
| 2 | Exon 6 F | AAAATTGCCCCAAAGCCCTG |
|  | Exon 6 R | GTGCTGGTGGAGAAGAGGAT |
| 3 | Exon 5 F | TGCTTCGGGCAGATAACACA |
|  | Myc.2 R | TCAGCTTCTGCTCTTGAGC |
| 4 | Exon 6 F | AAAATTGCCCCAAAGCCCTG |
|  | Myc.2 R | TCAGCTTCTGCTCTTGAGC |
